# Sex-specific expression and DNA methylation in a species with extreme sexual dimorphism and paternal genome elimination

**DOI:** 10.1101/2020.06.25.171488

**Authors:** Stevie A. Bain, Hollie Marshall, Laura Ross

## Abstract

Sexual dimorphism is exhibited in many species across the tree of life with many phenotypic differences mediated by differential expression and alternative splicing of genes present in both sexes. However, the mechanisms that regulate these sex-specific expression and splicing patterns remain poorly understood. The mealybug, Planococcus citri, displays extreme sexual dimorphism and exhibits an unusual instance of sex-specific genomic imprinting, Paternal Genome Elimination (PGE), in which the paternal chromosomes in males are highly condensed and eliminated from the sperm. P. citri also has no sex chromosomes and as such both sexual dimorphism and PGE are predicted to be under epigenetic control. We recently showed that P. citri females display a highly unusual DNA methylation profile for an insect species, with the presence of promoter methylation associated with lower levels of gene expression. In this study we therefore decided to explore genome-wide differences in DNA methylation between male and female P. citri using whole genome bisulfite sequencing. We have identified extreme differences in genome-wide levels and patterns between the sexes. Males display overall higher levels of DNA methylation which manifests as more uniform low-levels across the genome. Whereas females display more targeted high levels of methylation. We suggest these unique sex-specific differences are due to chromosomal differences caused by PGE and may be linked to possible ploidy compensation. Using RNA-Seq we identified extensive sex-specific gene expression and alternative splicing. We found cis-acting DNA methylation is not directly associated with differentially expressed or differentially spliced genes, indicating a broader role for chromosome-wide trans-acting DNA methylation in this species.

## Introduction

Sexual dimorphism is widespread across sexually-reproducing organisms. Males and females can differ dramatically in morphology, behaviour and physiology. Some of this dimorphism results from genetic adaptations that reside on sex chromosomes (Mank, 2009). However, many of these phenotypic differences are instead mediated by the differential expression of genes present in both sexes (Ellegren and Parsch, 2007). Sex-biased gene expression has been widely studied and varies amongst species, tissues and developmental stages (Grath and Parsch, 2016). However, the mechanisms that regulate these sex-specific expression patterns are often poorly understood.

DNA methylation is a well-characterised epigenetic modification that could facilitate such variation in expression (Grath and Parsch, 2016). DNA methylation is found throughout the genome of many organisms (Suzuki and Bird, 2008) and occurs most frequently at 5’-CG-3’ dinucleotides, known as CpG dinucleotides (Bird, 1986). In mammalian somatic tissue, 70-80% of all CpG sites are methylated (Feng *et al.*, 2010) and methylation at promoter regions can suppress gene transcription, leading to stable gene silencing (Bird, 2002). This is implicated in the regulation of sex-specific and sex-biased gene expression (examples include: Hall *et al.*, 2014; Maschietto *et al.*, 2017). In contrast, DNA methylation levels in arthropods are generally much sparser and vary across taxa (Thomas *et al.*, 2020). In most insects, DNA methylation is almost exclusively restricted to exons in a small subset of transcribed genes (Zemach *et al.*, 2010). The highest levels of global DNA methylation are found in hemimetabolous insects (e.g. 14% in Blattodea, Bewick *et al.*, 2017), while methylation is largely absent from holometabolous species (Provataris *et al.*, 2018; Lewis *et al.*, 2020). In insects, the role of DNA methylation in the regulation of gene expression remains inconclusive. However, studies show that DNA methylation is generally associated with elevated, stable gene expression (Foret *et al.*, 2009; Bonasio *et al.*, 2012; Wang *et al.*, 2013; Glastad *et al.*, 2016).

Despite evidence suggesting a relationship between DNA methylation and gene expression, few insect studies have directly explored sex-specific DNA methylation patterns and their association with sex-specific gene expression. In the jewel wasp, *Nasonia vitripennis*, 75% of expressed genes show sex-biased expression, however, DNA methylation patterns between the sexes are similar and do not explain gene expression patterns (Wang *et al.*, 2015). In contrast, a study in the peach aphid, *Myzus persicae*, in which 19% of genes exhibit sex-specific expression biases, reveals a correlation between sex-specific gene expression and sex-specific methylation, particularly for genes located on the sex chromosomes (Mathers *et al.*, 2019). Thus, the role of sex-specific patterns of methylation in regulating sex-biased gene expression in insects remains unclear.

The citrus mealybug, *Planococcus citri* (Hemiptera: Pseudococcidae), is uniquely suited for studying the functional role of DNA methylation in sex-specific gene expression. *P. citri* is a sexually reproducing species in which sexual dimorphism is extreme in morphology, life history and chromosome behaviour. Whilst the sexes are indistinguishable as nymphs, adult males and females are so morphologically distinct they could be mistaken as members of different species (Figure 1). Males undergo metamorphosis after the second instar and develop into winged adults (Sutherland, 1932). Females do not metamorphose, retain their larval appearance (neoteny), so remain wingless, and grow much larger than the males (Sutherland, 1932). In contrast to females, males do not feed after their second instar. Consequently, there is a large difference in lifespan between the sexes; with males only living up to 3 days after eclosion, while females can live several weeks after reaching sexual maturity (Nelson-Rees, 1960). Crucially, *P. citri* have no sex chromosomes meaning that males and females share the same genetic complement (Hughes-Schrader, 1948); therefore, the observed sexual dimorphism is solely a consequence of gene expression differences between the sexes.

**Figure 1:**
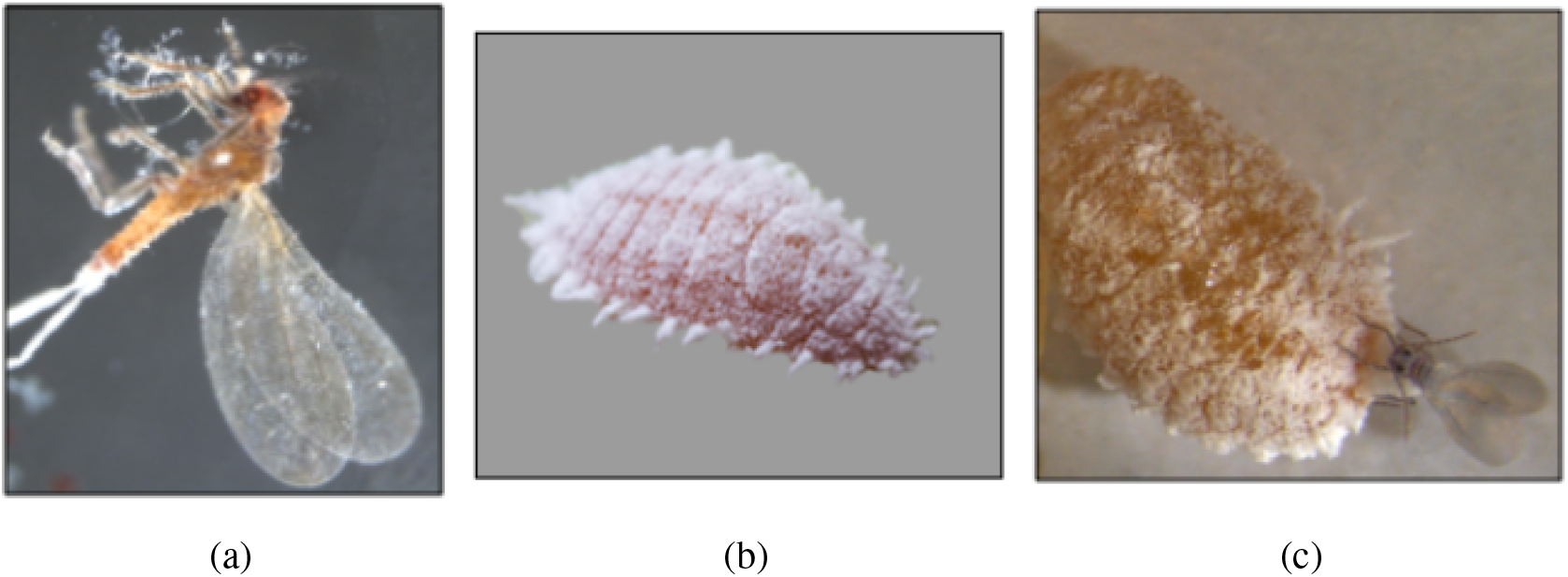
Extreme sexual dimorphism present in *Planococcus citri*. (a) Winged adult male, (b) neotenous adult female, (c) shows a male and female mating, where size difference between the sexes is apparent.

In addition to extreme sexual dimorphism, *P. citri* also has an unusual reproductive strategy, known as Paternal Genome Elimination (PGE). PGE is a genomic imprinting phenomenon found in thousands of insect species that involves the silencing and elimination of an entire haploid genome in a parent-of-origin specific manner. Under PGE, both sexes develop from fertilized eggs and initially possess a diploid euchromatic chromosome complement. However, males subsequently eliminate paternally-inherited chromosomes during spermatogenesis and only transmit maternally-inherited chromosomes to their offspring (Brown and Nelson-Rees, 1961). Furthermore, in *P. citri* males, paternally-inherited chromosomes are heterochromatinised in early development (Brown and Nur, 1964; Bongiorni *et al.*, 2001) and thus gene expression shows a maternal bias (de la Filia *et al.*, 2020). Females, on the other hand, do not undergo the process of PGE and both maternally and paternally-derived chromosomes remain euchromatic throughout development (Brown and Nur, 1964). Due to the haploidization of males, PGE is often referred to as a ‘pseudohaplodiploid’ system.

Furthermore, we have previously shown *P. citri* females have a unique pattern of whole genome DNA methylation that differs from that found in other arthropods (Lewis *et al.*, 2020). Whilst most arthropods have depleted levels of transposable element and promoter methylation, *P. citri* has independently evolved both (Lewis *et al.*, 2020). Interestingly, and similar to patterns shown in mammals, genes with low expression in *P. citri* have significantly higher promoter methylation than highly expressed genes (Lewis *et al.*, 2020). It is also suggested that DNA methylation may have a role in the recognition and silencing of paternally-derived chromosomes in males in the process of PGE (Bongiorni *et al.*, 1999; Buglia *et al.*, 1999). Supporting the idea that DNA methylation may be involved in sexual dimorphism and PGE in mealybugs and other scale insects, two recent studies have identified sex-biased expression of the DNA methyltransferase DNMT1 in adult *Phenacoccus solenopsis* (Omar *et al.*, 2020) and *Ericerus pela* (Yang *et al.*, 2015), with females showing considerably higher expression compared to males in both species.

In order to identify sex-specific patterns of gene expression and clarify the role of DNA methylation in this process, we analyse both male and female *P. citri* methylomes and transcriptomes. This is the first genome-wide analysis of sex-specific gene expression and DNA methylation in scale insects. Using RNA-seq and whole genome bisulfite sequencing (WGBS) we find clear differences in gene expression and methylation profiles between the sexes. However, we find no relationship between differentially expressed genes and differentially methylated genes, indicating that *cis*-acting DNA methylation is not the sole driver of sex-specific gene expression in adult *P. citri*.

## Materials and Methods

### Insect husbandry

Mealybug cultures used for this study were kept on sprouting potatoes in sealed plastic bottles at 25°C and 70% relative humidity. Under these conditions, *P. citri* has a generation time (time from oviposition until sexual maturity) of approximately 30 days. Experimental isofemale lines were reared in the laboratory under a sib-mating regime: in each generation, one mated female is taken per culture and transferred to a new container to give rise to the next generation. The *P. citri* line used (WYE 3-2) was obtained from the pest control company, WyeBugs in 2011, and had undergone 32 generations of sib-mating prior to this experiment. This high degree of inbreeding allows for precise mapping of Whole Genome Bisulfite-seq (WGBS) reads reducing mis-mapping caused by SNP variation. It also means we avoid contrasting methylation profiles caused by differences in the underlying genotype of individuals (epialleles).

We isolated virgin females after they became distinguishable from males (3rd-4th instar) and kept them in separate containers until sexual maturity (>35-days old). Males were isolated at the pupal stage and kept in separate containers until eclosion (~27 days). Insects were stored at −80°C until DNA and RNA extraction.

### RNA extraction and sequencing

We extracted RNA (3 biological replicates per sex, 60 males and 15 females per replicate) using TRIzol^®^ reagent (Thermo Fisher Scientific, USA) according to the manufacturer’s instructions and PureLink RNA purification kit (including DNase I digestion). Individual adult males are smaller than females; therefore, a higher number of males was required for each pooled sample. Samples were further purified with RNA Clean and Concentrator™-5. Quantity and quality of extracted genetic material was assessed using NanoDrop ND-1000 Spectrophotometer (Thermo Scientific, USA) and Qubit (Thermo Fisher Scientific, USA) assays. A260/A280 and A260/A230 ratios were calculated for all samples and only samples with A260/A280 of 1.7 - 2.0 and A260/A230 of >1.0 were processed. All RNA samples were sequenced by Edinburgh Genomics. Two of the samples (one male and one female) were sequenced on the Illumina HiSeq 4000 platform (75b paired-end reads). The remaining samples were sequenced on the Illumina NovaSeq S2 platform (50b paired-end reads).

### DNA extraction and bisulfite sequencing

We extracted genomic DNA from pools of 60 whole adult males and 15 whole virgin adult females using DNeasy Blood and Tissue kit (Qiagen, CA) and Promega DNA Clean and Prep Kit (Promega) in a custom DNA extraction protocol. Individual adult males are smaller than females; therefore, a higher number of males was required for each pooled sample. Five independent biological replicates were set up for each sex. DNA samples were cleaned and concentrated using Zymo DNA Clean and Concentrator Kit according to manufacturer’s instructions. DNA A260/A280 absorption ratios were measured with a NanoDrop ND-1000 Spectrophotometer (Thermo Scientific, USA) and concentrations were measured with a Qubit Fluorometer (Life Technologies, CA). Although five samples for each sex were prepared, two male samples had to be pooled in order to collect adequate DNA (500ng) for bisulfite conversion and library preparation. Therefore, there are only four male replicates.

Bisulfite conversion and library preparation was carried out by Beijing Genomics Institute (BGI). The bisulfite conversion rate is estimated based on non-methylated *Escherichia coli* lambda DNA (provided by BGI; isolated from a heat-inducible lysogenic *E. coli* W3110 strain. GenBank/EMBL accession numbers J02459, M17233, M24325, V00636, X00906), which was added at 1% to *P. citri* DNA samples. Sequencing of bisulfite libraries was carried out on an Illumina HiSeq4000 instrument to generate 150b paired-end reads.

### Differential expression and alternative splicing

Raw RNA-seq reads for each sample were trimmed for low quality bases and adapters using Fastp for paired-end reads (Chen et al., 2018). Fastp was used as it allows removal of poly-G tails from NovaSeq reads. We quantified gene-level expression for each sample using RSEM v1.2.31 (Li and Dewey, 2011) with STAR v2.5.2a (Dobin *et al.*, 2016) based on the *P. citri* reference genome and annotation (mealybug.org, version v0). Average expression and coefficient of variation was calculated per gene for individual male and female samples using FPKM (fragments per kilobase of transcript per million) values estimated by RSEM. Differentially expressed genes between the sexes were identified using EbSeq (Leng et al., 2013) based on gene-level expected counts produced by RSEM. A gene was considered differentially expressed if it had a fold-change >1.5 and a p-value < 0.05 after adjusting for multiple testing using the Benjamini-Hochberg procedure (Benjamini and Hochberg, 1995).

Alternatively spliced genes between sexes were identified using DEXSeq (Anders *et al.*, 2012) implemented by IsoformSwitchAnalyzeR (Vitting-Seerup and Sandelin, 2019). Briefly, this package implements a general linear model per gene which tests the relative proportion of expression of each exon per sex. This method accounts for within-sex gene expression differences and sex-specific gene expression differences. A gene was considered alternatively spliced if it had an absolute isoform usage difference of 10% and a p-value < 0.05 after adjusting for multiple testing using the Benjamini-Hochberg procedure (Benjamini and Hochberg, 1995).

### Genome-wide methylation patterns and differential methylation

Initial QC of Illumina reads was carried out using FastQC v.0.11.7 (Andrews, 2010). Quality and adapter trimming were carried out by BGI. *E. coli* and *P. citri* reference genomes (*P. citri* version v0, publicly available on mealybug.org) were converted to bisulfite format using Bismark Genome Preparation v0.19.0 (Krueger and Andrews, 2011). Illumina reads were first aligned to the converted unmethylated lambda *E. coli* control DNA sequence using Bismark v0.19.0 (Krueger and Andrews, 2011) to estimate the error rate of the C to T conversion. Bismark v0.19.0 and Bowtie2 were then used to align reads to the reference genome using standard parameters. The weighted methylation level of each genomic feature (*P. citri* v0 annotation, mealybug.org) was calculated as in Schultz *et al.* (2012). Briefly, this method accounts for the CpG density of a region by calculating the sum of all cytosine calls for every CpG position in a region (promoter/exon/gene etc.) divided by the total cytosine and thymine calls in the same region.

For differential methylation analysis between sexes coverage outliers (above the 99.9% percentile) and bases covered by < 10 reads were removed. Each CpG per sample was subjected to a binomial test to determine the methylation state, where the lambda conversion rate was used as the probability of success. Only CpGs which were determined as methylated in at least one sample were the tested via a logistic regression model, implemented using methylKit v1.10.0 (Akalin *et al.*, 2012), for differential methylation between the sexes. P-values were corrected for multiple testing using the Benjamini-Hochberg procedure (Benjamini and Hochberg, 1995). CpGs were considered differentially methylated if they had a q-value < 0.01 and a minimum methylation difference of 15%.

Promoter and exon regions were classed as differentially methylated if they contained at least three significant differentially methylated CpG sites and had a weighted methylation difference >15% across the entire region. Significant overlap of genes with promoter and exon differential methylation was determined using the hypergeometric test and visualised using the UpSetR package v1.4.0 (Lex *et al.*, 2016).

### Relationship of gene expression and DNA methylation

The relationship of promoter and exon methylation with gene expression and alternative splicing was assessed using custom R scripts. The mean FPKM and weighted methylation level was calculated across biological replicates for each sex. The presence of interaction effects in linear models was determined throughout using the *anova* function in R. Post-hoc testing of fixed factors was conducted using the *glht* function from the *multcomp* v1.4-12 R package with correction for multiple testing using the single-step method (Hothorn *et al.*, 2008). Correlations were calculated using Spearman’s rank correlation rho.

### Additional genome annotation

Promoter regions were defined as 2000bp upstream from each gene. We excluded promoters which overlap using BEDTools (Quinlan and Hall, 2010). Intergenic regions were determined as regions between the end of one gene and the beginning of the next gene’s promoter, excluding any annotated TEs. In order to determine possible sex-specific differences in transposable element (TE) methylation we annotated TEs within the *P. citri* genome. Following Lewis *et al.* (2020) we implemented RepeatModeller v.2.0 to create a model of TEs and then annotated these TE models with RepeatMasker v4.1.0 (http://www.repeatmasker.org). Differentially methylated CpGs were determined to originate from TEs if there was no genomic overlap with any other annotation, such as a gene body.

### Gene ontology enrichment

Gene ontology (GO) enrichment was carried out using the hypergeometric test with Benjamini-Hochberg correction for multiple testing (Benjamini and Hochberg, 1995), using the GOStats R package (Falcon and Gentleman, 2007). GO biological process terms were classed as over-represented if they had a q-value <0.05. REVIGO (Supek *et al.*, 2011) was used to visualise GO terms and obtain GO term descriptions. GO terms for genes with different levels of methylation were tested against a background of all genes. GO terms for genes which show female/male over expression were tested against a background of all genes identified in the RNA-Seq data. GO terms for genes which show extreme female/male over expression were tested against a background of all differentially expressed genes. GO terms for genes which show hypermethylation in either females/males were tested against a background of all genes identified in the WGBS data.

## Results

### Sex-biased gene expression and alternative splicing

All RNA-Seq samples generated between 66.9 million and 84.1 million paired-end reads with an average mapping rate of 87% (Supplementary 1.0.1). Genes showing different levels and patterns of sex-bias are likely subject to different evolutionary processes modulating their expression and sex-specificity (Wang, Werren and Clark, 2015). Therefore, in this study we distinguish three general categories of sex-biased genes. The first category contains sex-biased genes, defined as having >1.5-fold difference in expression between the sexes (q <0.05). The second contains extremely sex-biased genes, which are those that show >10-fold difference in expression between the sexes (q <0.05). The third category consists of sex-limited genes, i.e. those with some level of expression in one sex but no detectable expression in the other sex.

*P. citri* shows extreme sex-specific expression with many genes showing complete sex-limited expression (Fig.2a). We have identified a total of 10,548 significant genes with sex-biased expression between *P. citri* males and females (Fig.2b, Supplementary 1.0.2). This is 26.5% of the estimated 39,801 genes in the *P. citri* genome and 54.7% of all genes identified as expressed in at least one sex in the RNA-Seq data (n = 19,282). Of these sex-biased genes, 10,026 show moderate sex-biased expression (q <0.05 and >1.5 fold change) with significantly more showing female biased expression (5,270 compared to 4,756, chi-squared goodness of fit: X-squared = 26.351, df = 1, p <0.001). GO term enrichment analysis of sex-biased genes show that both female and male biased genes are enriched for core biological processes such as biosynthetic processing and carbohydrate metabolism (Supplementary 1.0.3). Additionally, female sex-biased genes are enriched for the GO term “*methylation*” (GO:0032259) and male sex-biased genes are enriched for “*chitin metabolic process*” (GO:0006030).

**Figure 2:**
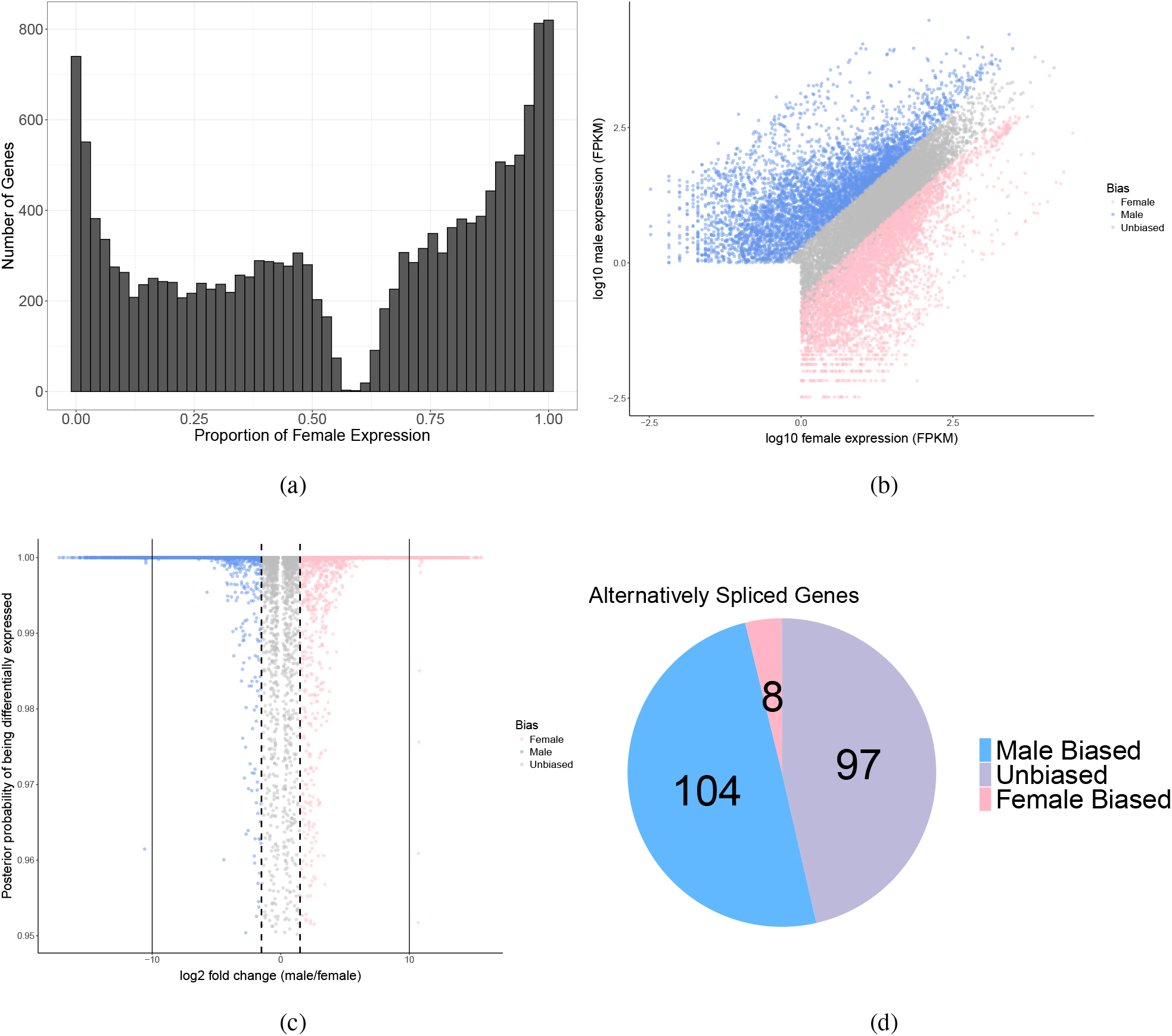
(a) Histogram of the proportion of female expression per gene for all genes present in the RNA-Seq data (n = 19,282). (b) Scatter graph of the log10 fragments per kilobase of transcript per million mapped reads (FPKM) for male and female samples. Each point is a gene. (c) The difference in the fold-change of genes plotted against the probability of being differentially expressed. Each point is a gene. (d) Pie chart showing the number of alternatively spliced genes which are also differentially expressed (male/female bias) or unbiased.

We also identify 168 extremely sex-biased genes (q <0.05 and >10 fold change, Supplementary 1.0.2), with the majority of these showing extreme male-biased expression (140 compared to 28, chi-squared goodness of fit: X-squared = 74.667, df = 1, p <0.001, Fig.2c). There were only three GO terms enriched for extremely biased male genes, these were “*system process*” (GO:0003008), “*sensory perception*” (GO:0007600) and “*sensory perception of smell*” (GO:0007608). Female *P. citri* are known to produce pheromones to attract males (Bierl-Leonhardt *et al.*, 1981), therefore it may be that these extremely male-biased genes are involved in pheromone response. There were no enriched GO terms for genes showing extreme expression bias in females.

Finally, we identify 354 sex-limited genes (q <0.05 and zero expression in one sex, Supplementary 1.0.2) in *P. citri*. Of these, significantly more are sex-limited to males compared to females (204 compared to 150, chi-squared goodness of fit: X-squared = 8.2373, df = 1, p = 0.01). GO terms enriched for female sex-limited genes include: “*growth*” (GO:0040007) and “*anatomical structure development*” (GO:0048856) amongst others (Supplementary 1.0.3). GO terms enriched for male sex-limited genes include: the same three GO terms mentioned above for extreme sex-biased genes as well as “*proteolysis*” (GO:0006508) and some other more general terms (Supplementary 1.0.3).

Next we searched for alternative splicing differences between the sexes. In the current genome annotation, (*P.citri* v0 mealybug.org), 93.13% of genes are annotated as single isoforms. After filtering out genes which also have low expression in both sexes (<10 FPKM), 1,235 genes were tested for alternative splicing. 209 genes were found to be significantly alternatively spliced between the sexes, consisting of 423 isoforms (q <0.05 and a minimum percentage difference of 25%, Supplementary 1.0.4). The GO terms enriched for alternatively spliced genes are varied, including some related to protein modification (Supplementary 1.0.5).

We next checked to see if any of the same genes show both sex-biased expression and sex-biased alternative splicing. We found that there was a significant overlap of alternatively spliced genes and genes with sex-specific expression bias (112/209), hypergeometric test, p <0.001). The majority of these genes (104/112) also show higher levels of male expression compared to higher female expression (chi-squared goodness of fit: X-squared = 82.286, df = 1, p <0.001, Fig.2d). There were no GO terms enriched for female bias and unbiased alternatively spliced genes compared to all alternatively spliced genes as a background. However, male biased alternatively spliced genes were enriched for metabolic processes and “*proteolysis*” (GO:0006508) (Supplementary 1.0.5).

The sex-determination system in *P. citri* is unknown and alternative splicing of the *doublesex* gene has been implicated in sex-determination in the vast majority of insect species (Wexler *et al.*, 2019). We therefore checked to see if any genes orthologous to the *Drosophila melanogaster doublesex* gene (as determined in: de la Filia *et al.*, 2020) were alternatively spliced. There were five genes in the current annotation (*P.citri* v0 mealybug.org) which are othlogous to *D. melanogaster doublesex* (*g1737*, *g2969*, *g11101*, *g11102* and *g36454*). We checked the list of differentially alternative spliced genes between sexes and none of these genes were differentially alternatively spliced, suggesting the method of sex-differentiation in this species is not via alternative splicing of *doublesex*. We also checked for sex-biased expression of these genes and only two were expressed, *g2969* and *g36454*, the former shows unbiased expression and the latter shows male-biased expression although overall expression levels are low. Finally, it is also worth noting *transformer*, a gene required for *doublesex* splicing (Wexler *et al.*, 2019) is not present in the *P. citri* genome.

### Sex-specific DNA methylation: genome-wide trends

Mapping rates for samples to the *P. citri* reference genome were 53.6% ± 2.8% (mean ± standard deviation). This equated to 24,220,848 ± 1,786,437 reads, which after deduplication gave an average coverage of 14.5X ± 1.1X (Supplementary 1.0.7). The bisulfite conversion efficiency across samples, calculated from the lambda spike, was 99.53% ± 0.05%. After correcting for this the single-site methylation level (Schultz *et al.*, 2012) in a non-CpG context was calculated as 0.05% ± 0.05% for females and 0.13% ± 0.05% for males. In a CpG context, females have significantly lower methylation levels compared to males, 7.8% ± 0.35% and 9.28% ± 0.26% respectively (Fig.3a, t-test: t = −7.17, df = 6.99, p <0.001). Additionally, using genome-wide CpG methylation levels males and females cluster separately, with females clustering much more tightly compared to males (Fig.3b). The diversity within male samples may be explained by a lower input of DNA during the library preparation process, resulting in possible sequencing bias.

**Figure 3:**
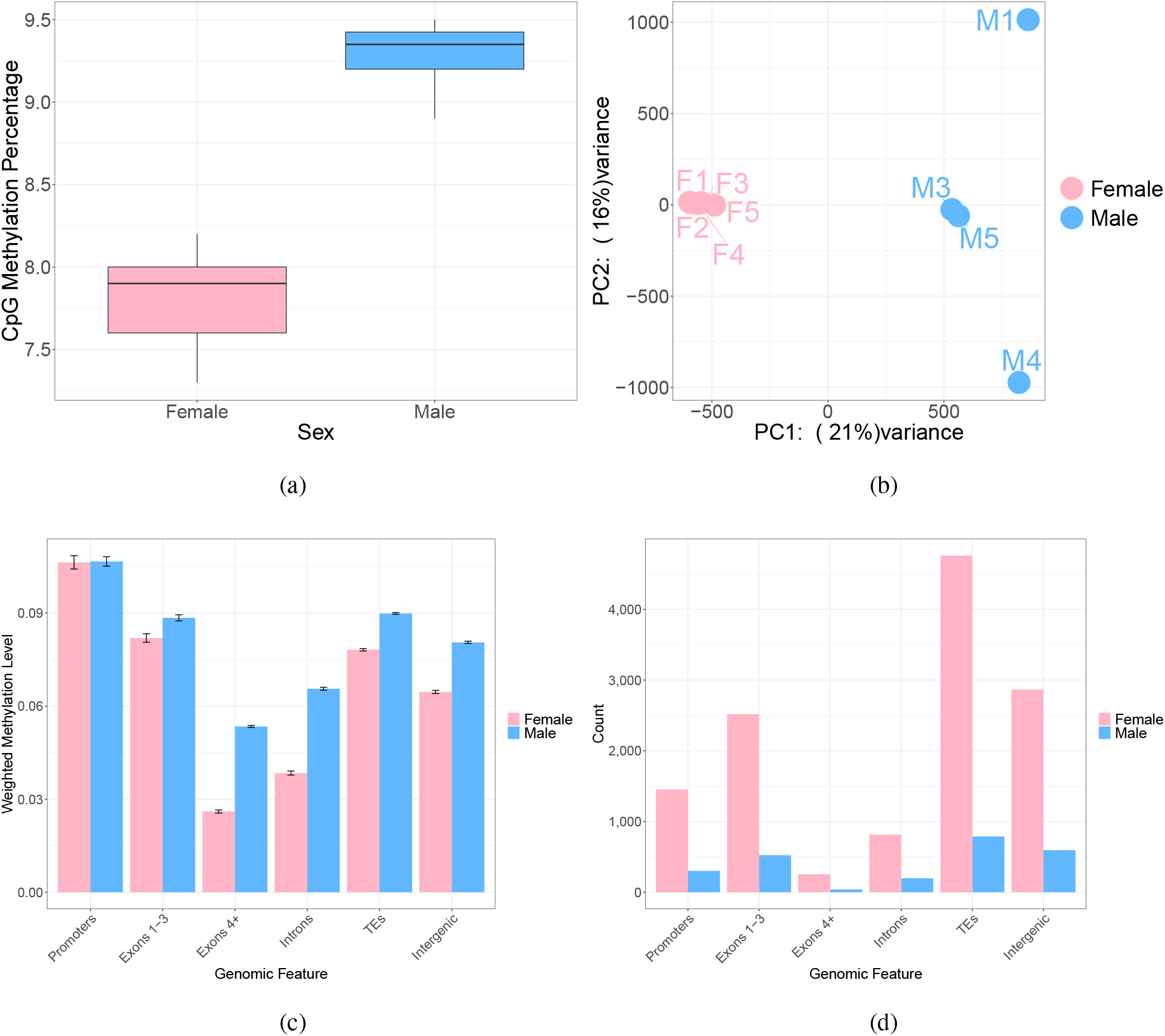
(a) Boxplot of the mean single-site methylation level in a CpG context for female and male replicates. (b) PCA plot generated by methylKit using per site CpG methylation levels. (c) The mean weighted methylation level for each genomic feature by sex, the error bars represent 95% confidence intervals of the mean. (d) Bar plot of the total number of genomic features which have a weighted methylation >0.7 in each sex.

Males and females also show significant differences in the distribution of CpG methylation levels across genomic features (two-way ANOVA, interaction between genomic feature and sex; F = 316.54, p <0.001, Fig.3c). As we have previously shown using the female data in Lewis *et al.* (2020) *P. citri* females have unusual patterns of DNA methylation compared to other insect species; we confirm that promoters, exons 1-3 and transposable elements (TEs) have significantly higher levels of DNA methylation compared to exons 4+ and introns (Fig.3c, Supplementary 1.0.8). We have found this is also the case for males, however, males show significantly higher levels of methylation than females in all features except for promoters (Supplementary 1.0.8). Additionally, we also found high levels of intergenic methylation in both sexes, which, along with promoter and TE methylation is highly unusual in insect species (Bewick *et al.*, 2019).

In order to determine the distribution of methylation levels across features, we binned features into four categories: highly methylated (>0.7), medium levels of methylation (0.3-0.7), lowly methylated (0-0.3) and no methylation, and plotted the number of features which fall into each bin per sex. We found that females show a more bimodal pattern of methylation and have significantly more features that show a weighted methylation level >0.7. For example, in the promoter region female n =1454 and male n =303 (Test of equal proportions, q <0.001, Fig.3d) and in exons 1-3 female n =1815 and male n =366 (Test of equal proportions, q <0.001, Fig.3d). Females also show significantly more genes with zero methylation in these features than males (Supplementary 2.0, Table S1.). This shows that the higher overall levels of genome methylation in males are driven by a larger number of lowly methylated features (Supplementary 2.0 Fig.S1 and S2), suggestive of uniform low levels of DNA methylation across the genome.

Due to the bimodal nature of female DNA methylation, we hypothesised that genes with different levels of methylation may be involved in different functions in males and females. For example, high levels of DNA methylation in insects has been associated with highly expressed housekeeping genes (Provataris *et al.*, 2018). Indeed, we found that genes with different levels of promoter and exon methylation are enriched for different functions. Highly methylated genes (>0.7 weighted methylation) in males and females are enriched for metabolic and core cellular processes (Supplementary 1.0.9) suggesting at least some highly methylated genes may be housekeeping genes. Genes with medium levels of methylation (0.3-0.7 weighted methylation) are also enriched for metabolic processes. Genes with low levels of methylation (0-0.3 weighted methylation) contain a large and general variety of terms. Unmethylated genes have enriched GO terms for protein-related processes. Additionally, three out of nine enriched GO terms for genes with no exon methylation in males are related to mRNA splice site selection (GO:0006376, GO:0000398, GO:0000375); these terms are not enriched for genes with no exon methylation in females. This suggests DNA methylation could play a functional role in alternative splicing in males.

### Sex-specific DNA methylation: gene-level

There were 3,660,906 CpG sites found in all replicates with a minimum coverage of 10X. 75.8% of these were classed as methylated in at least one sample by a binomial test and were then used for differential methylation analysis. A total of 182,985 CpGs were classed as differentially methylated between males and females (q <0.01 and a minimum percentage difference of 15%), which is around 5% of all CpGs in the genome. The majority of these sites are located in exons 1-3, promoters and TEs (Fig.4a). Exons 1-3 have the highest density of differentially methylated CpGs, followed by promoters and then TEs (Supplementary 2.0 Fig.S3)

**Figure 4:**
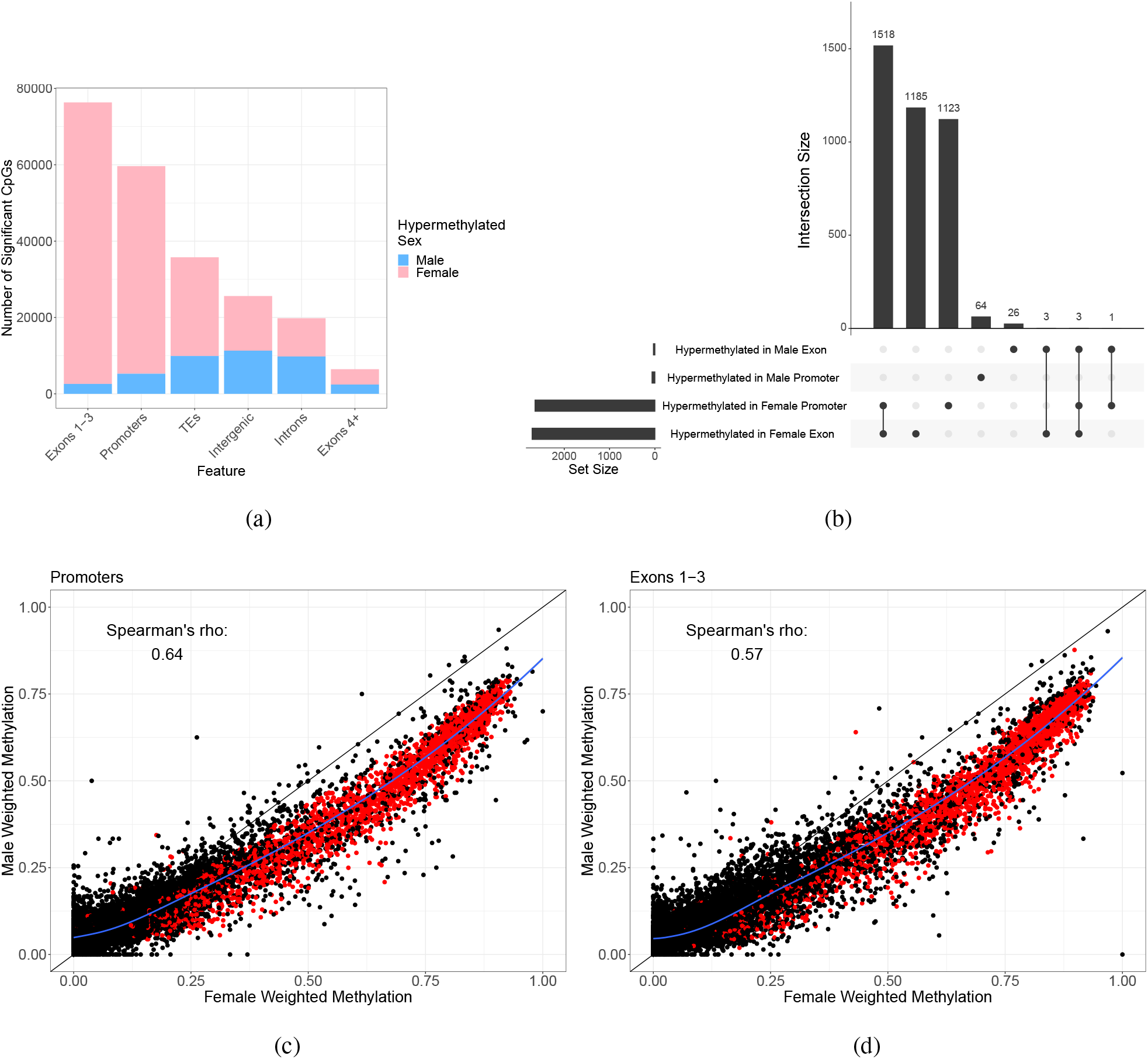
(a) Component bar plot showing the number of differentially methylated CpGs per genomic feature per sex. (b) UpSet plot showing the overlap between genes which are hypermethylated in either males or females. The set size indicates the total number of genes per group. The interaction size shows how many overlap or are unique to each group. Overlaps are shown by joined dots in the bottom panel, a single dot refers to the number of unique genes in the corresponding group. (c) and (d) scatter plots of the weighted methylation level of promoters and exons respectively. Each dot represents one promoter or one exon, the red dots are those which are significantly differentially methylated. The blue line represents a LOESS regression with the shaded grey area representing 95% confidence areas.

Due to the unusual occurrence of promoter methylation in *P. citri*, we investigated sex-specific differences in promoter methylation and exon 1-3 methylation, separately. We find 2,709 genes with a differentially methylated promoter (minimum three differentially methylated CpGs and a minimum overall weighted methylation difference of 15%) between males and females and 2,736 genes with differentially methylated exons 1-3 (Supplementary 1.1.0). A significantly higher number of genes with differential promoter methylation were hypermethylated in females compared to males (2,645 in females and 64 in males, chi-squared goodness of fit, X-squared = 2459, df = 1, p < 0.001, Fig.4b and 4c). This was also the case for genes with differential exon methylation, with 2,709 hypermethylated in females and 33 hypermethylated in males (chi-squared goodness of fit, X-squared = 2611.6, df = 1, p < 0.001, Fig.4b and 4d). In females, there is also a significant overlap of genes showing both hypermethylation of the promoter region and exons 1-3 (hypergeometric test, p <0.001, Fig.4b).

As males show mostly low-to-medium levels of methylation throughout the genome, we analysed the distribution of methylation levels for each feature determined as differentially methylated. We found that the average level of methylation in males for male hypermethylated promoters is 0.12 ± 0.07 (mean ± standard deviation) and for exons is 0.14 ± 0.12 (Supplementary 2.0, Fig.S4a and S4b), meaning the minimum 15% threshold difference applied translates into a small actual difference in methylation between males and females. Female hypermethylated sites were confirmed to show a full range of levels (Supplementary 2.0, Fig.S4a and S4b). The average level of female methylation for female hypermethylated promoters was 0.7 ± 0.18 and for exons was 0.73 ± 0.15. Whilst the differential methylation analysis conducted here is particularly stringent and in line with previous work on non-model insect species (e.g. Mathers *et al.*, 2019; Marshall *et al.*, 2019; Arsenault *et al.*, 2018), male hypermethylated sites should be interpreted with care.

In females, GO terms enriched for genes with hypermethylated promoters and exons were similar, mostly metabolic and DNA related processes such as, “*DNA integration*” (GO:0015074) and “*DNA replication*” (GO:0006260) (Supplementary 1.1.1). In males, only one GO term was enriched for genes with hypermethylated promoters, “*protein prenylation*” (GO:0018342). GO terms enriched for genes with hypermethylated exons in males were more diverse, including: “*response to pheromone*” (GO:0019236) (Supplementary 1.1.1).

### Relationship of DNA methylation and expression

Gene body DNA methylation is reported to positively correlate with gene expression in a number of insect species (Foret *et al.*, 2009; Bonasio *et al.*, 2012; Wang *et al.*, 2013; Glastad *et al.*, 2016; Marshall *et al.*, 2019). However, *P. citri* females show a negative relationship as higher methylation is correlated with lower gene expression (Lewis *et al.*, 2020). We explored this relationship further by examining both exon 1-3 and promoter methylation in males and females. On a single gene level, higher promoter methylation is significantly associated with lower gene expression (linear model: df = 63932, t = −10.44, p <0.001, Fig.5a and 5b). This is the case for both males and females as there is no interaction between sex and methylation level (two-way ANOVA: F_2,3_ = 0.265, p = 0.606). However, as there are few sites with high methylation in males (>0.75) this trend is curtailed. The same relationship is found between gene expression and methylation of exon 1-3 (Supplementary 2.0: Fig.S5a and S5b).

**Figure 5:**
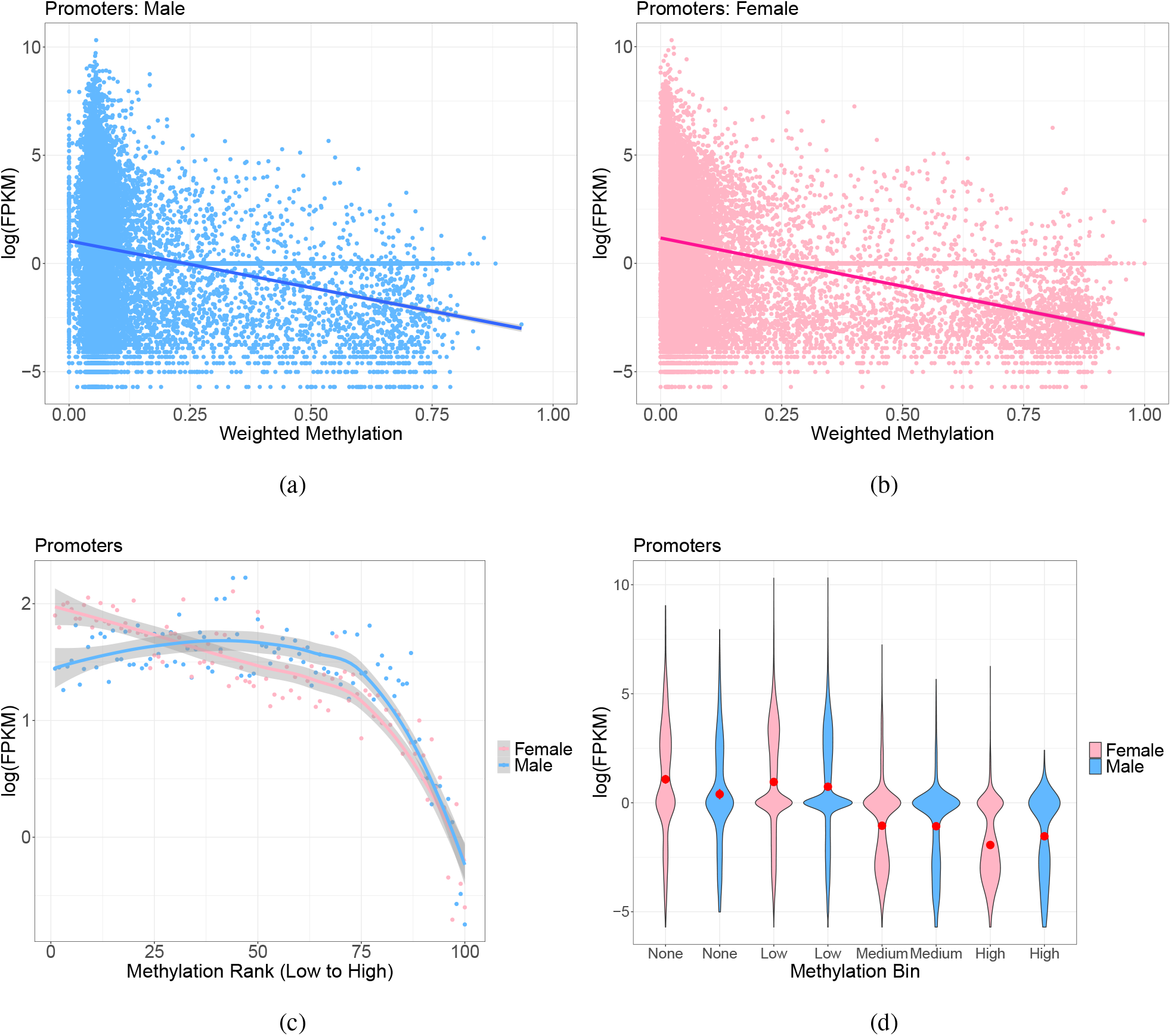
(a) and (b) scatter graphs of expression levels of every gene plotted against the mean weighted methylation level across replicates of each gene’s promoter region for males and females respectively. Each point represents one gene. The lines are fitted linear regression with the grey areas indicating 95% confidence intervals. (c) Genes were binned by mean weighted methylation level of the promoter region across replicates and the mean expression level of each bin as been plotted for males and females. The lines are LOESS regression lines with the grey areas indicating 95% confidence areas. (d) Violin plots showing the distribution of the data via a mirrored density plot, meaning the widest part of the plots represent the most genes. Weighted methylation level per promoter per sex, averaged across replicates, was binned into four categories, no methylation, low (>0–0.3), medium (0.3–0.7), and high (0.7–1). The red dot indicates the mean with 95% confidence intervals.

On a genome-wide scale the relationship between gene expression and methylation becomes more apparent (Fig.5c). Genes with no promoter methylation and low levels of promoter methylation have significantly higher expression levels compared to genes with medium and high promoter methylation (linear model: low methylation bin: df = 63930, t = 4.93, p <0.001, no methylation bin: df = 63930, t = 4.047, p <0.001, Fig.5d). Again, there is no interaction between sex and methylation bin (two-way ANOVA: F_4,7_ = 0.998, p = 0.392). The results for exon 1-3 methylation are similar, however, only the low methylation bin has significantly higher expression than genes with medium, high or no exon 1-3 methylation (Supplementary 2.0: Fig.S5c and S5d).

### Relationship of differential DNA methylation and differential expression

If DNA methylation is a causative driver of changes in gene expression we would expect that differentially methylated genes between sexes are also differentially expressed. Given that higher methylation is associated with lower expression in this species, we would also expect that down-regulation of gene expression is associated with higher methylation. However, on a single gene level, we found there is no clear relationship between the level of differential promoter methylation and the level of differential expression of the corresponding gene (Fig.6a). This is also the case for exon 1-3 methylation (Supplementary 2; Fig.S6a).

**Figure 6:**
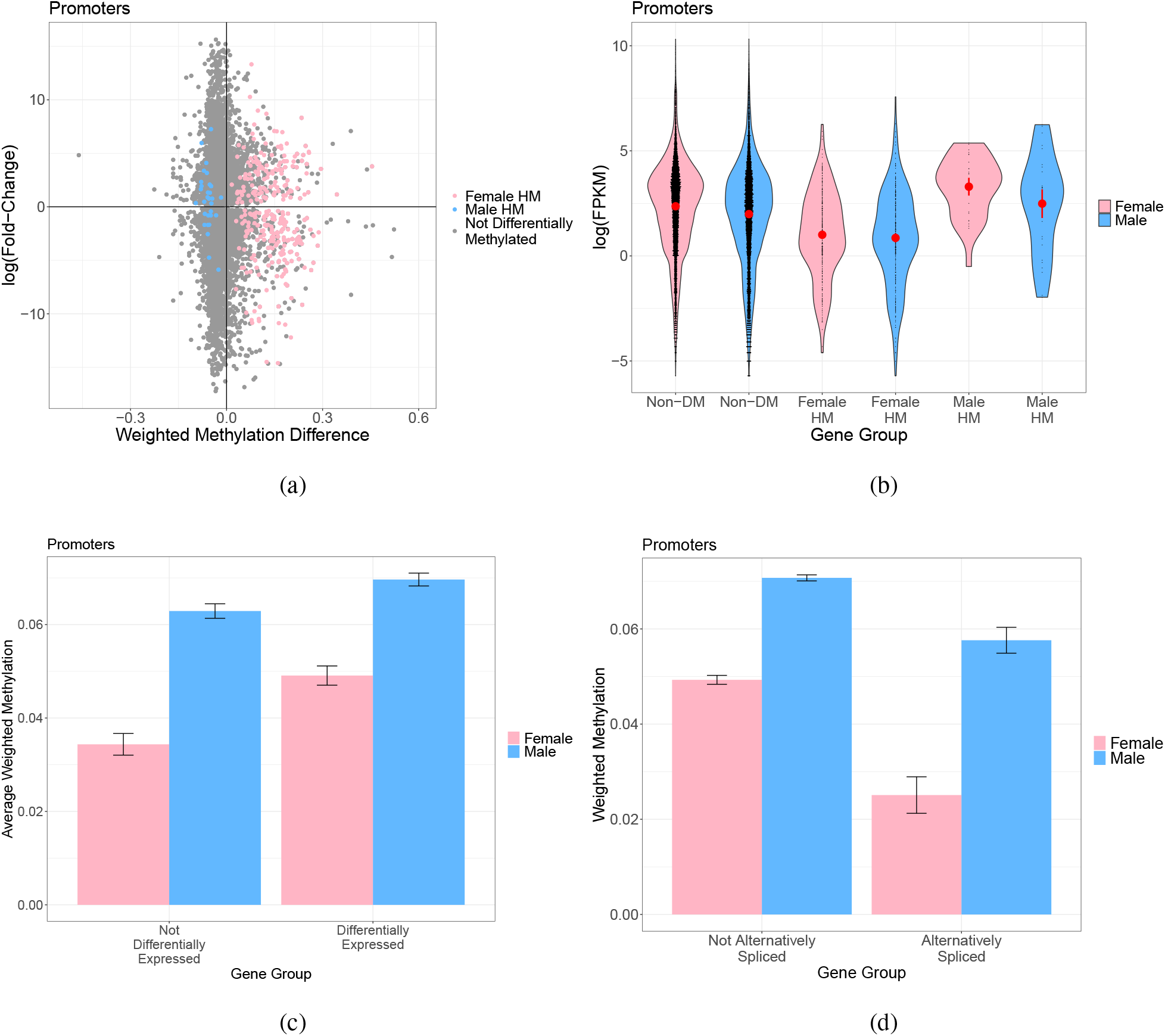
(a) Scatter plot of the weighted methylation difference between sexes (mean female weighted methylation minus mean male weighted methylation) for promoters plotted against the log fold-change in gene expression. A log fold-change greater than zero represents over expression in females. Each point represents a single gene. Blue points are genes which have significant male promoter hypermethylation and pink points are genes which have significant female promoter hypermethylation. (b) Violin plot of the expression levels of genes which are not differentially methylated between sexes (Non-DM) or which are hypermethylated (HM) in either females or males. Each black point is a gene. The red dot represents the mean with 95% confidence intervals. (c) Bar plot of the mean weighted methylation level of the promoter regions for differentially expressed genes and unbiased genes. Error bars represent 95% confidence intervals of the mean. (d) Bar plot of the mean weighted methylation level of promoter regions for genes which are alternatively spliced or not. Error bars represent 95% confidence intervals of the mean.

Additionally, genes that are hypermethylated in female promoter regions are enriched for genes that show significant expression bias in both females (overlapping genes = 113, hypergeometric test with bonferroni correction, p <0.001) and males (overlapping genes = 92, hypergeometric test with bonferroni correction, p = 0.024). Genes that are hypermethylated in female exons 1-3 are enriched for genes with just female biased expression, the opposite of our prediction (overlapping genes = 138, hypergeometric test with bonferroni correction, p <0.001, Supplementary 2: Table S2). Finally, male hypermethylated genes are not significantly enriched for any genes which show sex-biased expression but male hypermethylated promoters are enriched for unbiased genes (overlapping genes = 14, hypergeometric test with bonferroni correction, p = 0.021, Supplementary 2: Table S2). Therefore, whilst genome-wide higher methylation is correlated with lower expression, this trend is not replicated on a single gene basis, indicating *cis*-acting DNA methylation does not drive differences in gene expression between the sexes.

We next explored general expression levels of differentially methylated genes. We found that genes with hypermethylated promoters in females show significantly lower levels of expression compared to those with non-differentially methylated promoters (Tukey post-hoc: t = −2.756, p <0.05, Fig.6b). Interestingly, the expression levels of these female hypermethylated genes are similar in both sexes (two-way ANOVA for the interaction of sex and differentially methylated category: F_3,5_ = 0.013, p = 0.987 Fig.6b). The expression levels of genes which have hypermethylated promoters in males appear similar to genes with non-differentially methylated promoters and are not significantly different to those with female hypermethylated promoters (Tukey post-hoc: t = 0.642, p = 0.782, Fig.6b). The same relationships are observed when genes with differentially methylated exons are assessed (Supplementary 2: Fig.S6b).

We then assessed the overall methylation levels of differentially expressed genes. We found the average promoter methylation level of differentially expressed genes is higher than for unbiased genes in both sexes (linear model: df = 27375, t = −10.136, p <0.001, Fig.6c). The same differences are also observed with exon 1-3 methylation (Supplementary 2: Fig.S6c). We then checked to see if a specific set of sex-biased genes, such as those which are sex-limited, drive this overall methylation difference observed. We found no significant difference in methylation between biased, extremely biased and sex-limited categories. We also found the pattern of higher male promoter/exon 1-3 methylation compared to female methylation is the same in most cases (Supplementary 2: Fig.S7a and S7b). Finally, it is worth noting we also found annotated genes which were not present in the RNA-Seq data set had considerably higher methylation levels in males and females compared to genes which were expressed in either sex (Supplementary 2: Fig.S8a and S8b).

### Relationship of DNA methylation and alternative splicing

Exonic DNA methylation has been associated with alternative splicing in some insect species (Bonasio *et al.*, 2012; Li-Byarlay *et al.*, 2013; Marshall *et al.*, 2019). Therefore, we tested for a relationship between DNA methylation and sex-specific alternative splicing in *P. citri*. We found that unlike differentially expressed genes, the promoter methylation levels of alternatively spliced genes are lower than non-alternatively spliced genes (linear model: df = 27612, t = −3.772, p <0.001). The same pattern is also observed with exon 1-3 methylation (Supplementary 2: Fig.S6d). Additionally, alternatively spliced genes which also show sex-specific expression bias do not significantly differ in their promoter or exon 1-3 methylation levels compared to alternatively spliced genes which show unbiased expression (Supplementary 2: Fig.S7c and S7d).

We also then checked to see if alternatively spliced genes were also differentially methylated between sexes. We found only one significant overlap of genes which are both alternatively spliced and differentially methylated (Supplementary 2: Table S3), a single gene was common between alternatively spliced genes which show male expression bias and genes with male promoter hypermethylation (hypergeometric test with bonferroni correction, p = 0.034). However, it is likely this overlap is significant due to the small gene lists rather than due to biological significance.

## Discussion

In this study, we investigated the relationship between sex-specific gene expression and DNA methylation in the mealybug, *Planococcus citri*, a species with extreme sexual dimorphism and genomic imprinting (PGE). Our major findings include: the identification of vastly different genome-wide methylation profiles between the sexes, high levels of intergenic methylation - especially in males, and no relationship between differentially expressed genes and differentially methylated genes, indicating *cis*-acting DNA methylation does not regulate sex-specific differences in adult gene expression.

We hypothesise that the DNA methylation patterns we observe can be explained by several mechanisms acting simultaneously: 1) the higher and more even distribution of methylation across the male genome could be a cause or consequence of the heterochromatinization of the paternal genome in males, 2) the regulation of a subset of mostly non-sexually dimorphic genes through promoter/exon methylation in both sexes, 3) the hypermethylation of certain promoters and exons reducing expression in females, possibly to balance expression level between the sexes as a mechanism of ploidy compensation.

### PGE may explain uniform DNA methylation in males

We have identified extreme sex-specific differences in DNA methylation across the genome of *P. citri*. Most notably, overall higher genome-wide methylation levels in males manifest as low, uniform levels across the genome in comparison to a more targeted bimodal pattern of DNA methylation in females. To our knowledge, this type of sex-specific pattern has not been reported in any other species to date. We have also confirmed promoter methylation in both sexes, which is highly unusual in insects (Lewis *et al.*, 2020). We hypothesise this pattern, along with the identification of intergenic DNA methylation, is a result of the unusual reproductive strategy employed by this species, paternal genome elimination. Males with PGE have approximately half of their genome in a heterochromatic state (Hughes-Schrader, 1948; Brown and Nur, 1964; Bongiorni and Prantera, 2003; de la Filia *et al.*, 2020). In mammals and plants, DNA methylation is associated with the formation of heterochromatin (Suzuki and Bird, 2008). Previous research has found DNA methylation differences between the paternal and maternal chromosomes in mealybug species, although studies do not agree upon which chromosome set shows higher levels of DNA methylation (Bongiorni *et al.*, 1999; Buglia *et al.*, 1999; Mohan and Chandra, 2005). It is therefore likely the differences in the pattern of DNA methylation between the sexes may be driven by the condensed paternal chromosomes in males. Future work utilising reciprocal crosses to identify parent-of-origin DNA methylation at base-pair resolution throughout the genome would further clarify the role of DNA methylation in chromosome imprinting in this species.

Whilst differences in DNA methylation have been associated with the different parental chromosomes, it is the modifications of histones which have been directly linked to the formation of heterochromatin in *P. citri* (reviewed in Prantera and Bongiorni, 2011). Most recently Bain (2019) showed that both the H3K9me3-HP1 and H3K27me3-PRC2 heterochromatin pathways are involved in the condensation of the paternal chromosomes in males. Additionally, non-CpG methylation is also thought to exist in mealybugs in a CpA and CpT context (Deobagkar *et al.*, 1982) and the genes coding for the necessary enzymatic machinery for these modifications have recently been identified in the mealybug *Maconellicoccus hirsutus* (Kohli *et al.*, 2020). Although we did not find methylation levels above 0.2% in any non-CpG context (Supplementary 1.0.7). These studies suggest PGE is likely mediated by multiple interactions between a variety of epigenetic mechanisms within the genome.

### DNA methylation in females may be involved in ploidy compensation

Another striking pattern we observe is the hypermethylation of single CpG sites in female (compared to male) promoters and exons. Overall hypermethylation in females suggests DNA methylation in males and females may serve different functions. We hypothesise that one function of hypermethylation in females could be to act as a mechanism of ploidy compensation, as due to paternal chromosome silencing, most genes show haploid expression in males (de la Filia *et al.*, 2020). There is evidence for possible ploidy compensation via DNA methylation in other insects. Elevated DNA methylation levels in haploid males of the fire ant, *Solenopsis invicta*, are suggested to be indicative of regulatory pressures associated with the single-copy state of haploid loci (Glastad *et al.*, 2014). The aphid *Myzus persicae*, also shows male hypermethylation on the X chromosome which appears as a single copy in males (Mathers *et al.*, 2019). Although, it should be noted female aphids show much higher DNA methylation in the autosomes which are diploid in both sexes. However, it known in mammals that DNA methylation serves multiple functions in the genome (e.g. Edwards *et al.*, 2017) and this has also been suggested to be the case in insects with the function of DNA methylation potentially changing depending on the genomic context (Glastad *et al.*, 2018). In the examples noted above higher methylation has been identified in the sex/chromosome which is in the haploid state. DNA methylation in these species is associated with elevated, stable gene expression (Mathers *et al.*, 2019; Hunt *et al.*, 2013), suggesting methylation in these examples may serve to increase expression levels to compensate for single gene copies. We find a negative relationship between DNA methylation and gene expression in *P. citri*, suggesting higher methylation in females may serves to decrease expression of certain genes to mirror the haploid expression levels of males. This is further supported by our finding that female hypermethylated genes show overall similar expression levels in both females and males. To test this idea the expression levels of non-sex-biased genes from each parental chromosome set in both males and females should be assessed. Balanced expression levels would suggest some form of ploidy compensation.

We find no consistent overlap between differentially methylated genes and differentially expressed genes. This suggests that *cis*-acting DNA methylation is not regulating sex-specific gene expression. However, if DNA methylation does indeed play a role in ploidy compensation we would expect to see no overlap with differentially expressed genes. These findings further support the idea that DNA methylation is involved in chromosome-wide processes, such as paternal chromosome condensation in males and possibly ploidy compensation in females. Indeed, a recent RNAi study which knocked down *DNMT1* in the mealybug *Phenacoccus solenopsis*, found phenotypic changes in males and females, with females changing colour and losing their waxy coating and males displaying wing abnormalities (Omar *et al.*, 2019). This supports this idea that DNA methylation is involved in the generation of sex-differences in mealybugs. However, another RNAi study in the Hemipteran, *Oncopeltus fasciatus*, revealed that depletion of DNA methylation did not result in changes in gene or transposable element expression but did lead to aberrant egg production and follicle development (Bewick *et al.*, 2019). Thus, suggesting a functional role for DNA methylation that is independent to specific gene expression. It is also worth noting that previous work in insects has found conflicting evidence for the role of DNA methylation in differential gene expression. Wang *et al.* (2015) found no correlation between methylation and sex-specific expression in a species of Nasonia. Whereas, Mathers *et al.* (2019) found differentially methylated genes between aphid sexes were enriched for differentially expressed genes. Future experimental validation, such as in Omar *et al.* (2019) and Bewick *et al.* (2019), exploring specifically the functional role of methylation in regulating gene expression in diverse insect species is sorely needed.

### Sex-specific expression and splicing mirror extreme sexual dimorphism

In addition to our key findings above we have also identified sex-specific gene expression and alternative splicing. *P. citri* have no sex chromosomes meaning that males and females share the same genetic complement (Hughes-Schrader, 1948). Thus, the observed sexual dimorphism exhibited must be a consequence of differences in gene expression and splicing between the sexes. Indeed, we found that 54% of genes show sex-biased expression, including a subset of genes that are extremely sex-biased and sex-limited. We found that both male- and female-biased genes are involved in core biological processes. Sex-limited genes are likely important in the phenotypic sex differences observed in *P. citri*, including sensory related male-limited genes that may be involved in mate recognition through pheromones (Bierl-Leonhardt *et al.*, 1981). Nasonia males also show extreme sex-biased expression of pheromone genes (Wang *et al.*, 2015). The large number of differentially expressed genes we have identified reflects the extreme sexual dimorphism shown in this species (Fig.1).

We also identified differentially alternatively spliced genes between the sexes and found a significant number of these show male-biased expression. Genome-wide sex-specific alternative splicing has also been identified in aphids (Grantham and Brisson, 2018) and other insects (e.g. Glastad *et al.*, 2016; Price *et al.*, 2018; Rago *et al.*, 2020). Specifically, Grantham and Brisson (2018) found that differentially expressed and alternatively spliced genes had similar GO term enrichment and they suggest both mechanisms serve to independently generate phenotypic differences between the sexes. Given the significant overlap of differentially expressed and differentially alternatively spliced genes we have found here, it may be that *P. citri* utilises expression regulation and alternative splicing of many of the same pathways to generate phenotypic sex differences. Additionally, Gibilisco *et al.* (2016) have shown male and female Drosophila utilise alternative splicing differently - males increase diversity in their gene expression profiles by expressing more genes and females express less genes but use more alternative transcripts. In *P. citri*, we found generally more female-biased genes compared to male-biased genes but more male-biased alternatively spliced genes, showing that *P. citri* sexes also employ different mechanisms to generate sex-specific phenotypes.

Surprisingly, we did not find any genes orthologous to the Drosophila *doublesex* gene to be alternatively spliced. Alternative splicing of *doublesex* is ubiquitous in holometabolous insects, whereas male-biased expression rather than alternative splicing has been detected in some crustaceans (Kato *et al.*, 2011; Li *et al.*, 2018) and a mite (Pomerantz and Hoy, 2015), indicating male-biased expression was likely the ancestral mode of *doublesex* sex-differentiation (Wexler *et al.*, 2019). Recently, Wexler *et al.* (2019) explored the role of *doublesex* orthologs in three hemimetbolous insect species and concluded the splicing method of sexual differentiation has evolved within the hemipteran order. One of the identified *doublesex* orthologs (*g36454*) in *P. citri* shows male biased expression indicating a possible ancestral function. Although it is worth noting expression levels of this gene are low in both sexes. Improved functional annotation of the current genome build may uncover isoforms not currently identified. Additionally, future work is needed to experimentally validate the role *g36454* may have in sex differentiation.

### Future Considerations

It is important to bear in mind that the differences we describe in this study are found in adult whole body samples and thus do not capture expression and DNA methylation biases between tissues and developmental stages, which are known to vary greatly (Harrison *et al.*, 2015; Grath and Parsch, 2016). Recently both sex-specific and developmental stage specific expression has been identified in other mealybug species: *Phenacoccus solenopsis* (Omar *et al.*, 2019), *Planococcus kraunhiae* (Muramatsu *et al.*, 2020) and *Maconellicoccus hirsutus* (Kohli *et al.*, 2020). With Kohli *et al.* (2020) identifying sex-specific expression of numerous epigenetic regulators, including the genes *SMYDA-4* and *SDS3* which are up-regulated in males and *SMYD5* and *nucleoplasmin* which are up-regulated in females. These genes are thought to be involved in heterochromatin formation via the methylation of various histones (Kohli *et al.*, 2020). The presence of histone marks is known to differ between sexes in mealybugs, with Ferraro *et al.* (2001) identifying higher histone acetylation in the paternal chromosome of *P. citri* males. The presence of such differences in adults may contribute to the extreme sexual dimorphism exhibited by mealybugs. In order to further understand the role of sex-specific expression, DNA methylation and other epigenetic modifications in *P. citri*, RNA-seq, ChIP-Seq/CUT&Tag and WGBS of specific tissues and developmental stages are needed.

## Conclusions

Overall, this study has shown striking differences in the DNA methylome of male and female *P. citri*, unlike any previously described sex-specific differences in insects. It is likely these differences are due to the unusual reproductive strategy of this species, paternal genome elimination. Based on our key finding of a lack of direct association between differential DNA methylation and differential gene expression, paired with recent findings by de la Filia *et al.* (2020) that show males display mostly haploid gene expression, we hypothesise DNA methylation may play a *trans*-acting role in ploidy compensation in this species, although this is speculation and requires experimental testing. Finally, we have identified a large number of differentially expressed genes between sexes mirroring the extreme sexual-dimorphism exhibited in this species and we have found no evidence for sex-specific alternative splicing of *doublesex* orthologs in *P. citri*. In addition to these key findings this study lays the groundwork for future research exploring the role of DNA methylation in genomic imprinting in insects as well as experimental validation studies to identify the interactions between multiple epigenomic mechanisms which may lead to such extreme sexual dimorphism and paternal genome elimination in this species.

## Acknowledgements

We thank Peter Sarkies for valuable advice and discussion. This work was supported by the NERC grant: NE/K009516/1, the Royal Society Grant: RG160842 and a Wellcome Trust - Institutional Strategic Support Fund awarded to L.R. S.A.B. was supported by the BBSRC Eastbio DTP. H.M. was supported by the ERC starting grant ‘PGErepro’ awarded to L.R.

## Data Accessibility

Data have been deposited in GenBank under NCBI BioProject: PRJNA610765. All code is available at: http://github.com/RossLab/Sex-Specific_Methylation_P.citri.

## Author Contributions

L.R. conceived the study. S.A.B. cultured the insects and conducted all lab work. S.A.B. carried out the differential expression analysis. H.M. carried out all other analyses with contribution from S.A.B. All authors wrote and reviewed the manuscript.

