## Supplementary 2 for "Sex-specific expression and DNA methylation in a species with extreme sexual dimorphism and paternal genome elimination"

S. A. Bain [1], H. Marshall [1] & L. Ross [1]

1) Institute of Evolutionary Biology, University of Edinburgh, UK.

### 2.0 Additional figures and tables

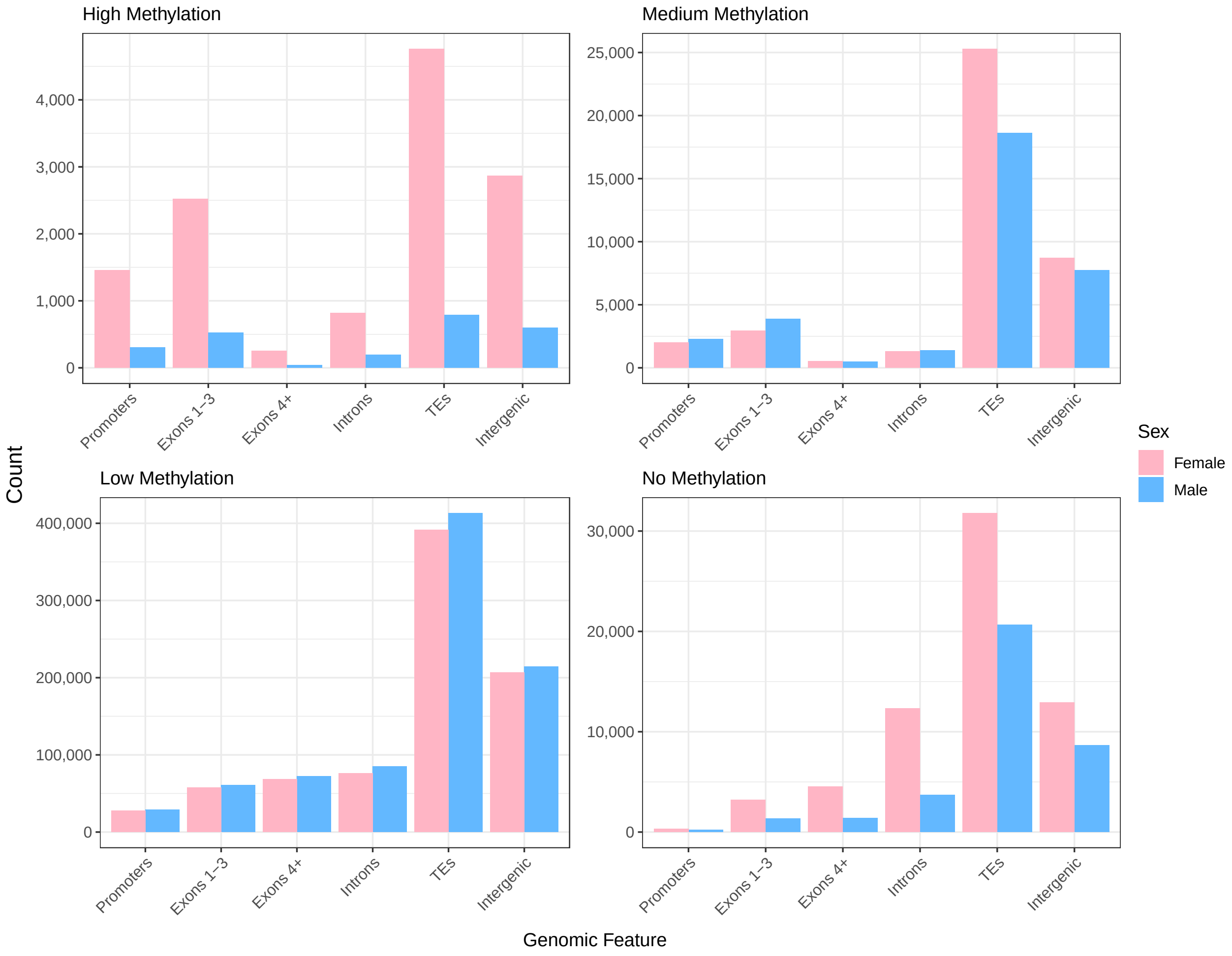


Figure S1: Bar plots of the total number of genomic features categorised by weighted methylation level for males and females. High methylation is a weighted methylation level >0.7, medium is 0.3-0.7, low is 0-0.3 and no methylation is equal to zero.

| Feature and methylation level | Male n | Female n | p-value | q-value |
|---|---|---|---|---|
| Promoters >0.7 | 303 | 1454 | $2.288\text{x}10^{-170}$ | $1.601\text{x}10^{-169}$ |
| Promoters 0.3-0.7 | 2283 | 1991 | $4.046\text{x}10^{-06}$ | $4.046\text{x}10^{-06}$ |
| Promoters 0-0.3 | 28949 | 27973 | $7.320\text{x}10^{-37}$ | $3.660\text{x}10^{-36}$ |
| Promoters 0 | 220 | 340 | $4.395\text{x}10^{-07}$ | $8.790\text{x}10^{-07}$ |
| Exons 1-3 >0.7 | 366 | 1815 | $3.722\text{x}10^{-218}$ | $2.977\text{x}10^{-217}$ |
| Exons 1-3 0.3-0.7 | 2851 | 2221 | $3.951\text{x}10^{-20}$ | $1.580\text{x}10^{-19}$ |
| Exons 1-3 0-0.3 | 29861 | 28862 | $8.261\text{x}10^{-51}$ | $4.957\text{x}10^{-50}$ |
| Exons 1-3 0 | 176 | 357 | $4.894\text{x}10^{-15}$ | $1.468\text{x}10^{-14}$ |

Table S1: Results of the test of equal proportions for the number of genes which have promoter/exon 1-3 methylation levels of different categories. Methylation level is the weighted methylation level. P-values were corrected for multiple testing using the Benjamini-Hochberg method.

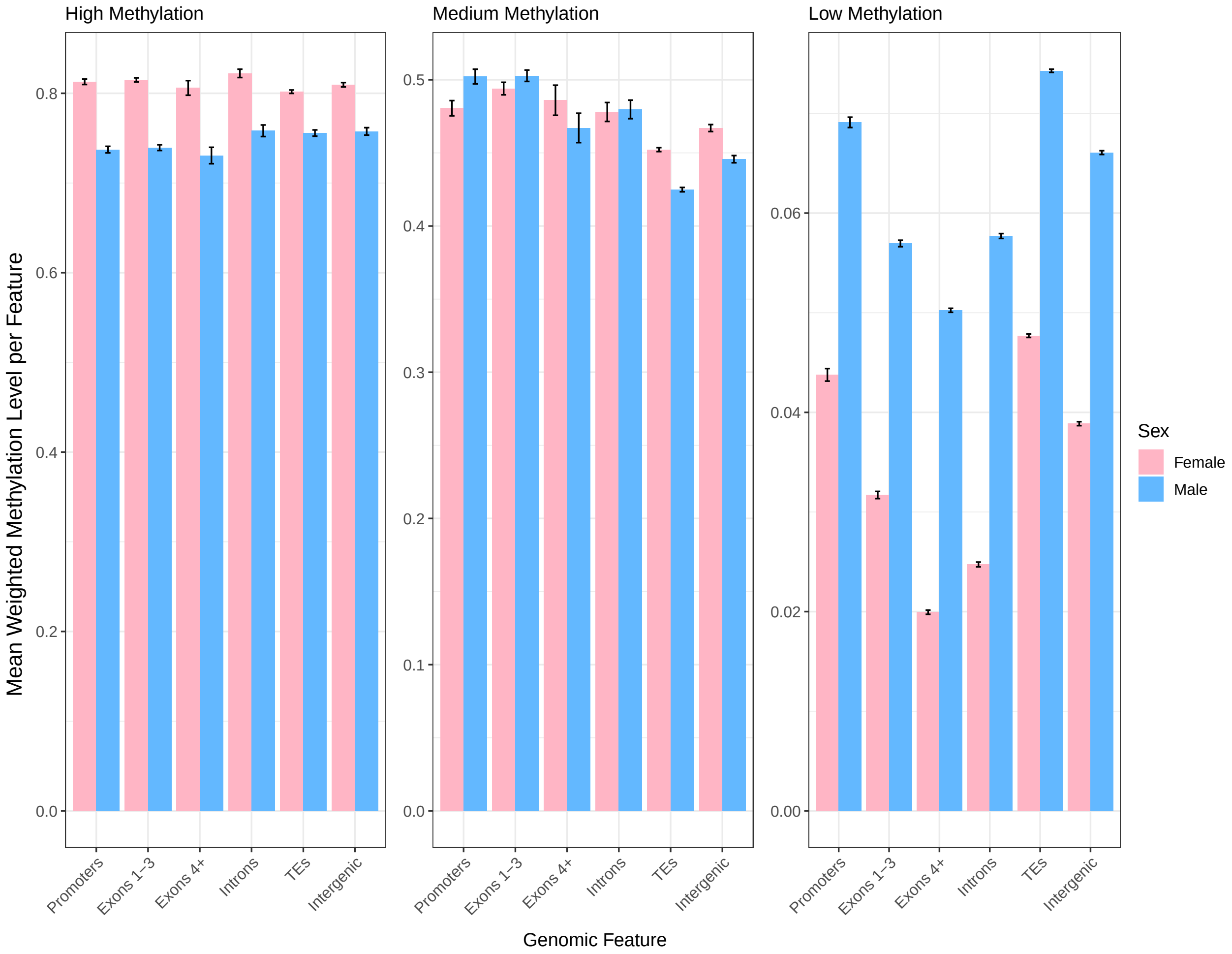


Figure S2: The mean weighted methylation level for each genomic feature by sex for the different categories of methylation. High methylation is a weighted methylation level >0.7, medium is 0.3-0.7 and low is 0-0.3. The error bars represent 95% confidence intervals of the mean.

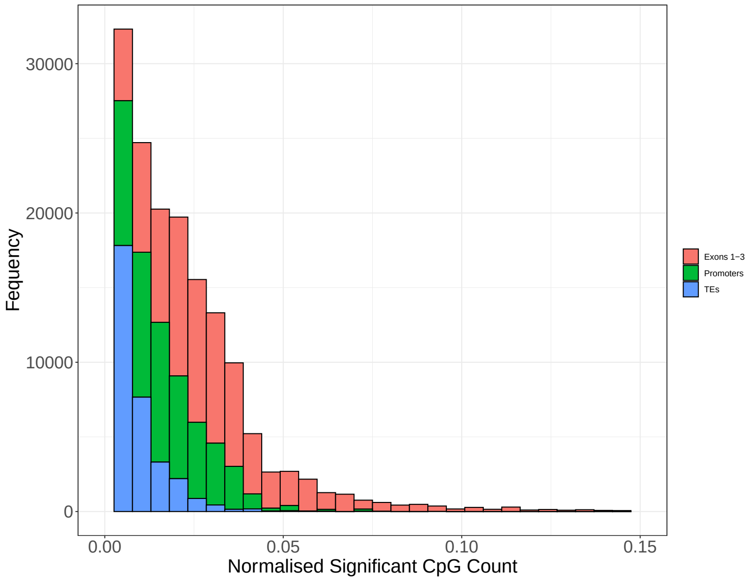


Figure S3: Histogram showing the number of differentially methylated CpGs between males and females per genomic feature, normalised by the length of each feature.

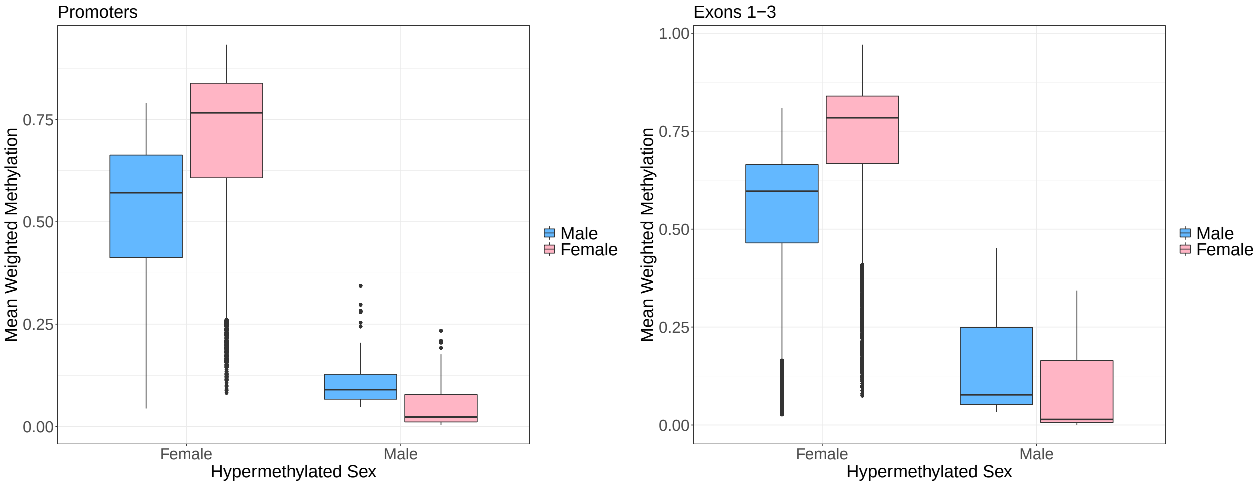


(a) (b)

Figure S4: Box plots of the weighted methylation levels of promoters (a) and exons 1-3 (b) for features which have been determined as differentially methylated (minimum three differentially methylated CpGs and a minimum overall weighted methylation difference of 15%) between sexes.

On a single gene level higher exon 1-3 methylation is significantly associated with lower gene expression (linear model effect of methylation of expression: df = 66943, t = -11.83, p <0.001, effect of sex on expression: df = 2, t = -0.40, p = 0.689, two-way ANOVA for the effect of an interaction between sex and methylation on expression: $F_{2,3} = 1.064$, p = 0.302).

On a genome-wide scale, genes with low exon 1-3 methylation have significantly higher expression than genes with medium, high or no exon 1-3 methylation(linear model: low methylation bin: df = 66941, t = 5.77, p <0.001, Fig. S5d). Again there was no effect of sex (linear model: df = 2, t = -0.741, p = 0.458) or an interaction between sex and exon 1-3 methylation (two-way ANOVA, $F_{4,7} = 0.116$, p = 0.95).

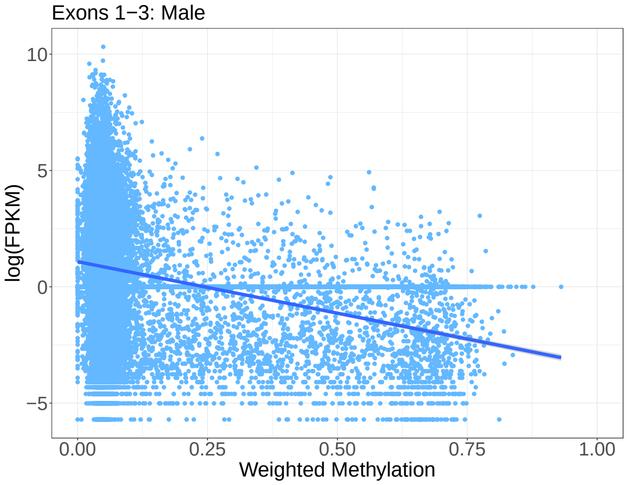


(a)

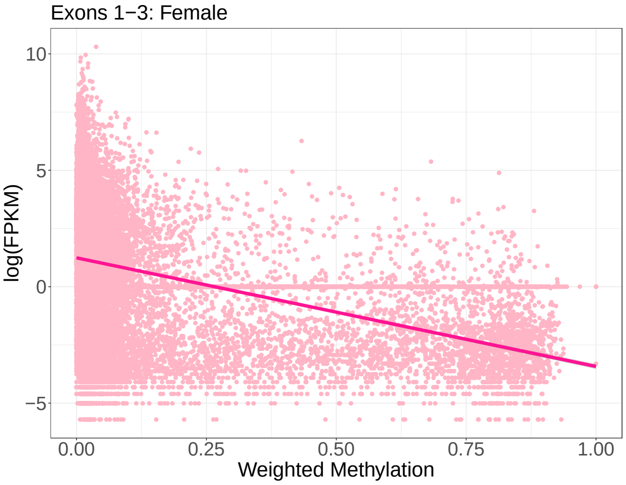


(b)

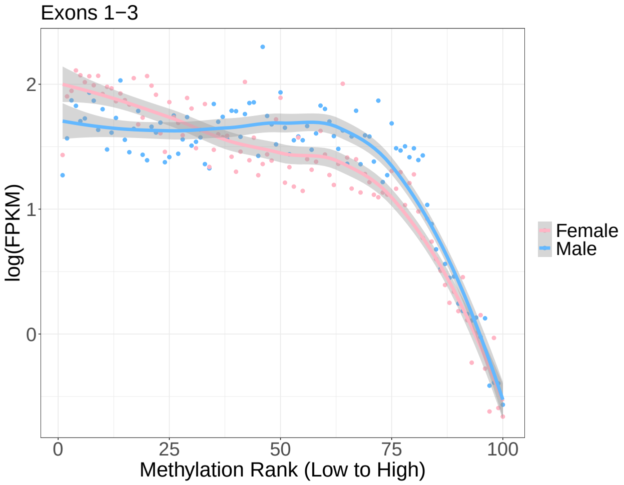


(c)

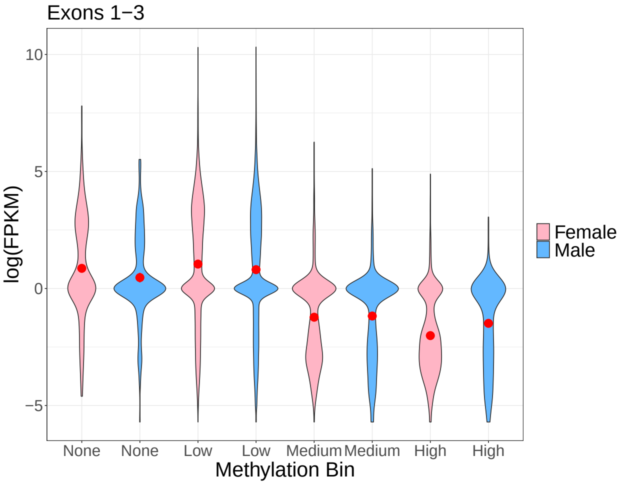


(d)

Figure S5: (a) and (b) scatter graphs of expression levels of every gene plotted against the mean weighted methylation level across replicates of each exon 1-3 region for males and females respectively. Each point represents one gene. The lines are fitted linear regression with the grey areas indicating 95% confidence intervals. (c) genes were binned by mean weighted methylation level across replicates and the mean expression level of each bin as been plotted for males and females. The lines are LOESS regression lines with the grey areas indicating 95% confidence areas. (d) Violin plots showing the distribution of the data via a mirrored density plot, meaning the widest part of the plots represent the most genes. Weighted methylation level per gene per exon 1-3, averaged across replicates, was binned into four categories, no methylation, low (0–0.3), medium (0.3–0.7), and high (0.7–1). The red dot indicates the mean with 95% confidence intervals.

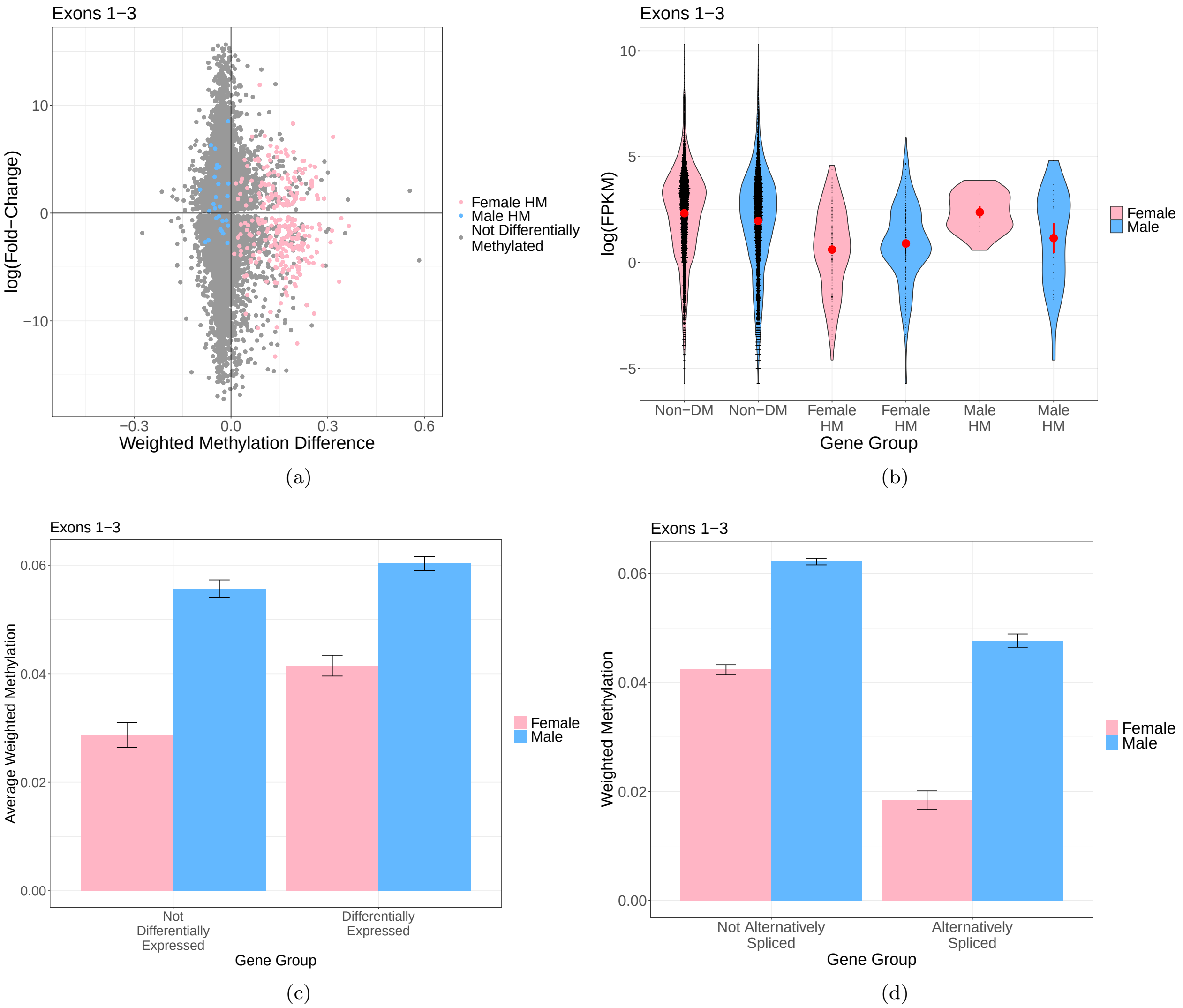


Figure S6: (a) Scatter plot of the weighted methylation difference between sexes (mean female weighted methylation minus mean male weighted methylation) for exons 1-3 plotted against the log fold-change in gene expression. A log fold-change greater than zero represents over expression in females. Each point represents a single gene. Blue points are genes which have significant male exon hypermethylation and pink points are genes which have significant female exon hypermethylation. (b) Violin plot of the expression levels of genes which are not differentially methylated between sexes (Non-DM) or which are hypermethylated (HM) in either females or males. Each black point is a gene. The red dot represents the mean with 95% confidence intervals. (c) Bar plot of the mean weighted methylation level of exons 1-3 for genes which are differentially expressed or not. Error bars represent 95% confidence intervals of the mean. (d) Bar plot of the mean weighted methylation level of exons 1-3 for genes which are alternatively spliced or not. Error bars represent 95% confidence intervals of the mean.

Table S2: The number of overlapping genes (n) which are both differentially methylated and show expression differences between sexes. Significant overlap was determined using the hypergeometric test with p-values corrected for multiple testing using the bonferroni method. HM refers to hypermethylation.

| | **Promoters** | | | | **Exons 1-3** | | | |
|---|---|---|---|---|---|---|---|---|
| | **Female HM** | | **Male HM** | | **Female HM** | | **Male HM** | |
| **Gene Expression Category** | **n** | **p-value** | **n** | **p-value** | **n** | **p-value** | **n** | **p-value** |
| Female Biased | 113 | <0.001*** | 6 | 1 | 138 | <0.001*** | 5 | 1 |
| Male Biased | 92 | 0.024* | 10 | 1 | 74 | 1 | 11 | 0.451 |
| Female Extreme Biased | 9 | 1 | 0 | N/A | 4 | 1 | 0 | N/A |
| Male Extreme Biased | 2 | 1 | 0 | N/A | 1 | 1 | 0 | N/A |
| Female Limited | 1 | 1 | 0 | N/A | 1 | 1 | 0 | N/A |
| Male Limited | 7 | 0.313 | 0 | N/A | 2 | 1 | 0 | N/A |
| Unbiased | 77 | 1 | 14 | 0.021* | 73 | 1 | 11 | 0.216 |

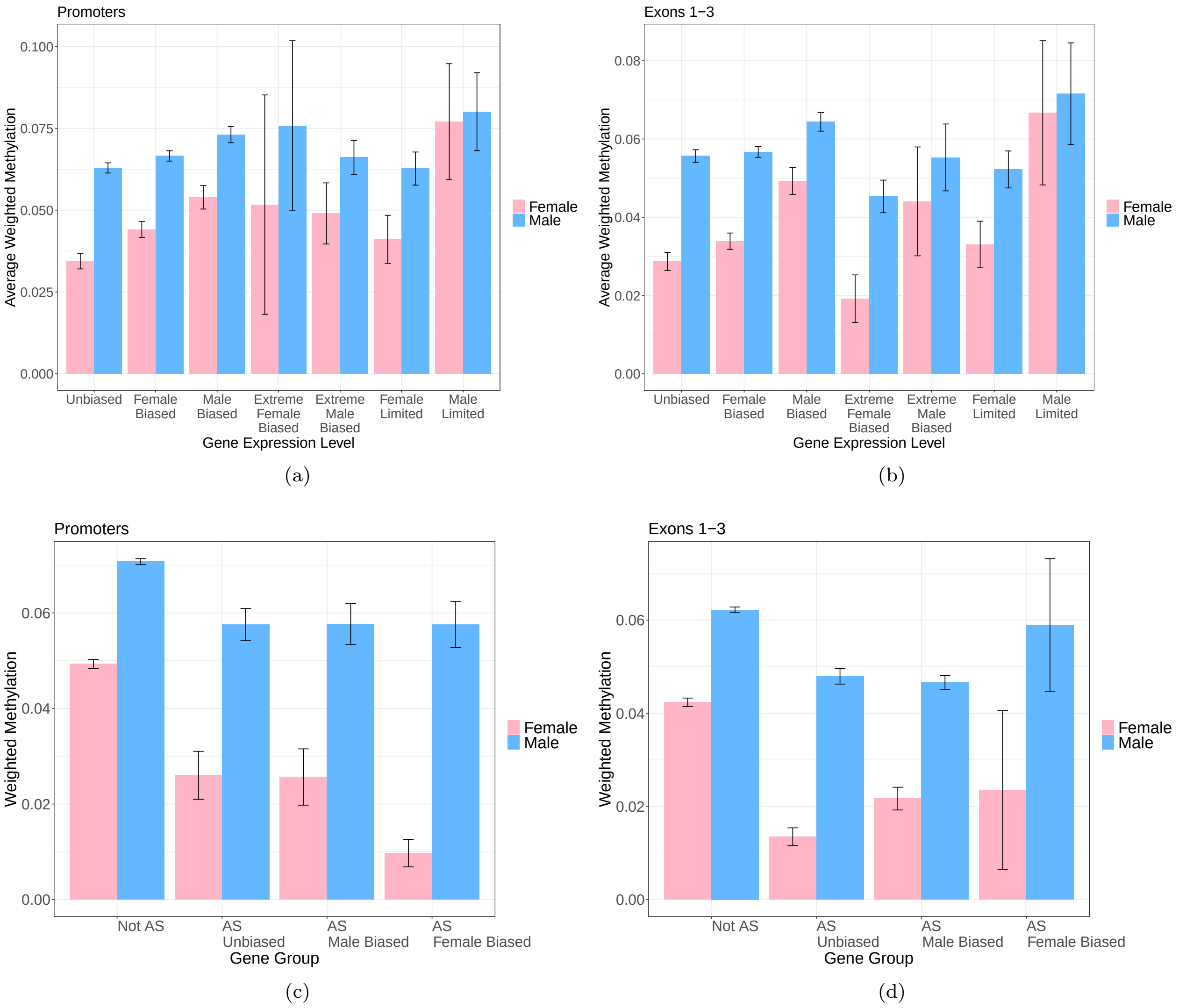


Figure S7: Bar plot of the mean weighted methylation level of (a) promoters and (b) exons 1-3 for genes which are differentially expressed or not. Differentially expressed genes are broken down into sex-biased, extremely sex-biased and sex-limited. Error bars represent standard error. (c) and (d) the same graph but for alternatively spliced genes broken down into those with significant male/female biased genes expression or unbiased gene expression.

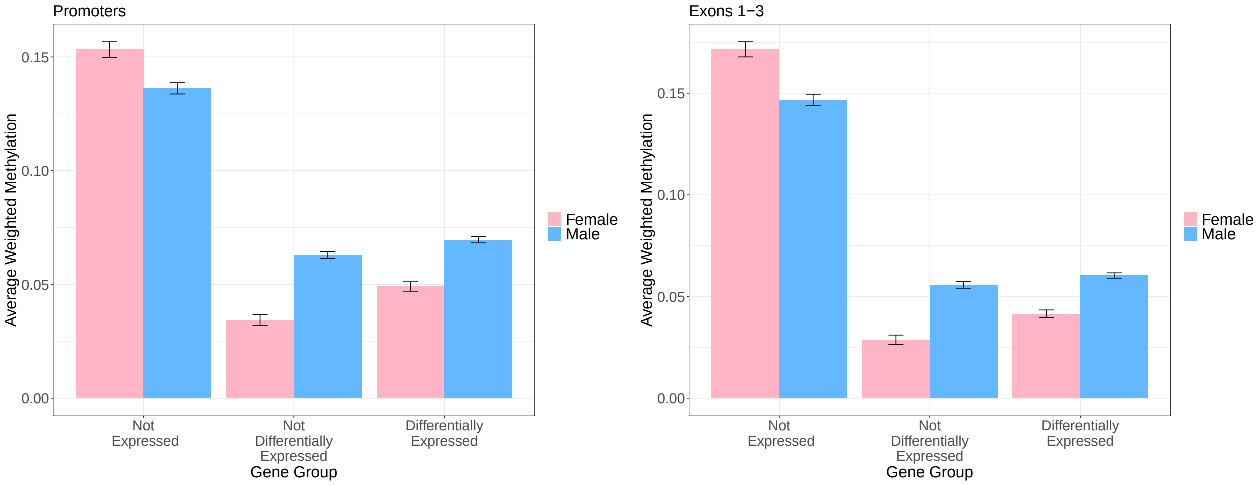


(a) (b)

Figure S8: Bar plot of the mean weighted methylation level of annotated genes which are not present in the RNA-seq as well as genes present in the RNA-seq which are differentially expressed or not, for (a) promoter methylation and (b) exon 1-3 methylation. Error bars represent 95% confidence intervals of the mean.

Table S3: The number of overlapping genes (n) which are both differentially methylated and show alternative splicing between sexes. Significant overlap was determined using the hypergeometric test with bonferroni corrected p-values. HM refers to hypermethylation.

| | **Promoters** | | | | **Exons 1-3** | | | |
|---|---|---|---|---|---|---|---|---|
| | **Female HM** | | **Male HM** | | **Female HM** | | **Male HM** | |
| **Alternative Splice Category** | **n** | **q-value** | **n** | **q-value** | **n** | **q-value** | **n** | **q-value** |
| Female Biased | 0 | N/A | 0 | N/A | 0 | N/A | 0 | N/A |
| Male Biased | 1 | 1 | 1 | 0.034* | 0 | N/A | 0 | N/A |
| Unbiased | 1 | 1 | 0 | N/A | 0 | N/A | 1 | 0.063 |
